# A novel inference of the fundamental biodiversity number for multiple immigration-limited communities

**DOI:** 10.1101/003137

**Authors:** Champak Beeravolu Reddy, Pierre Couteron, François Munoz

## Abstract

Neutral community theory postulates a fundamental quantity, θ, which reflects the species diversity on a regional scale. While the recent genealogical formulation of community dynamics has considerably enhanced quantitative neutral ecology, its inferential aspects have remained computationally prohibitive. Here, we make use of a generalized version of the original two-level hierarchical framework in order to define a novel estimator for *θ*;, which proves to be computationally efficient and robust when tested on a wide range of simulated neutral communities. Estimating *θ*; from field data is also illustrated using two tropical forest datasets consisting of spatially separated permanent field plots. Preliminary results also reveal that our inferred regional diversity parameter based on community dynamics may be linked to widely used ordination techniques in ecology. This paper essentially paves the way for future work dealing with the parameter inference of neutral communities with respect to their spatial scale and structure.

## Introduction

In his now well known contribution, Hubbell (2001) introduced a neutral theory of biodiversity, building upon the mainland-island model borrowed from the theory of island biogeography (MacArthur & Wilson 1967). His theory basically describes a model of interacting communities where the slow and regional scale dynamics such as speciation and extinction occur on a metacommunity level and the relatively faster local demographic events such as the birth and death of individuals are the mainstay of the local community dynamics. Hubbell (2001) further imagined equilibrium situations for these two entities which are dictated by the parameter *θ*(the fundamental biodiversity number) for the metacommunity dynamics and an immigration parameter *I* (Etienne & Olff 2004) for the local community dynamics. The interaction between these entities is modelled through the arrival of immigrant individuals from the regional pool of species (i.e. the metacommunity) towards a single or multiple local communities. It can be further interpreted as a measure of the isolation that a local community undergoes from the regional species pool due to some form of limited dispersal or more exactly immigration-limitation (Beeravolu *et al.* 2009). Up till now, much of the interest on neutral models in ecology has been either based on this two-level spatially implicit hierarchical framework (Hubbell 2001; Vallade & Houchmandzadeh 2003; Etienne 2005) or on spatially explicit models which do not make any such hierarchical distinctions (Chave & Leigh 2002; Zillio *et al.* 2005; O’Dwyer & Green 2009). In addition to model structure, details of the metacommunity processes such as speciation have been explored (Haegeman & Etienne 2008; Kopp 2010) though a consensus neutral approach is still lacking in community ecology (Gravel *et al.* 2006) and work is still in progress (Haegeman & Loreau 2011).

One explanation for the discord among ecologists, especially on the practical relevance of the two-level spatially implicit neutral model (hereafter denoted 2L-SINM), has been the issue of neutral parameter inference. Most of the current methods consist of either fitting a species abundance distribution (Purves & Pacala 2005; Dornelas *et al.* 2006; McGill *et al.* 2006) or finding quantitative point estimates of the neutral parameters (Etienne *et al.* 2006; Munoz *et al.* 2007). In the former, the general binning process of the species information into abundance classes, among other concerns (McGill *et al.* 2007), has raised several issues (Gray *et al.* 2006). At the same time the reliability of point estimates also remains doubtful (Leigh 2007), particularly as *I* and *θ* are known to be “hyperbolically correlated” when estimating them simultaneously (Beeravolu *et al. Submitted manuscript*; Etienne *et al.* 2006; Munoz *et al.* 2007; Beeravolu *et al.* 2009; Jabot & Chave 2009). Also, a basic contention underlying all these critiques has been the inherent insufficiency of the abundance information to fully elucidate ecological and evolutionary processes that neutral models aim to combine (Harte 2003).

In this paper, we build upon a recently discussed neutral framework which resembles a decoupling of scales (Levin 1992) and further enhances the spatial structure of the 2L-SINM. This is accomplished by introducing an intermediate level of regional process which generalizes the classical 2L-SINM (Hubbell 2001) into a 3L-SINM (Munoz *et al.* 2008) and enables the relaxation of the speciation-drift (or speciation-extinction) equilibrium assumption on the pool of available immigrant individuals (Beeravolu *et al.* 2009). In a previous paper, Munoz *et al.* (2008) introduced the 3L-SINM and used the steady state results to improve on the independent estimation of the neutral immigration parameter *I*, whereas we use similar approaches to establish an independent inference of *θ* for the first time.

## Methodological background

### The hierarchical SINM

Under the original 2L-SINM model, local communities are defined as panmictic patches of species assemblages sufficiently isolated from each other so as not to receive an immigrant individual directly from another local community (Fig. 1). Besides, from a regional perspective, local communities are presented as immigration-limited samples of the same metacommunity and subject to a top-down arrival of immigrants depending upon the immigration probability *m* (which is the scaled version of *I* on the unit scale). One possible generalization of the 2L-SINM can then be defined as a regional scale common to all local communities and whose species composition is allowed to differ considerably from that of the metacommunity, *sensu* Hubbell (2001, pg. 122), which is supposed to be panmictic and at speciation-drift equilibrium (Munoz *et al.* 2008). In more practical terms, immigrant species originating from a common source pool may very well correspond to a particular sub-region of the larger hypothetical metacommunity. For instance, let a mountain range be characterized by a coherent biogeographical history and represent a large-scale metacommunity. If a deep gorge runs through this range at some point, for all practical purposes, we can consider a part of the range to be under some sort of isolation from immigrants coming from the rest of the metacommunity, especially for the case of sessile biota such as plants even though their overall floristic composition remains specific to the mountain biotope.

**Figure 1.**
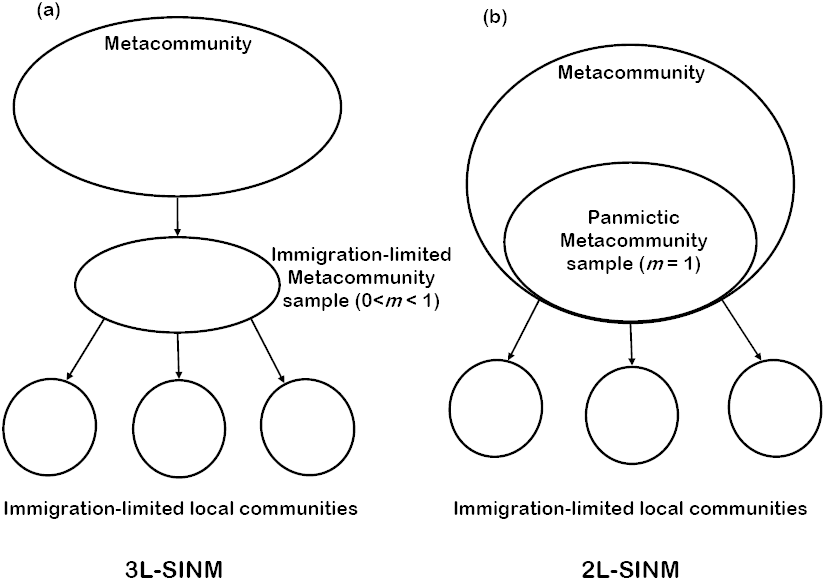
The hierarchical coupling of scale oriented processes in spatially implicit neutral models (SINM). Processes such as speciation and extinction of species exist in the metacommunity and the relatively faster demographic processes take place at the local community level. For Hubbell’s (2001) two level model (2L-SINM) an ideal panmictic metacommunity sample can be considered to be the source pool of multiple local communities. Without loss of generality, we can assume that the pooled species composition of the dispersal- limited local community samples is itself a immigration-limited metacommunity sample (the 3L-SINM) corresponding to some sub-region of the same which gives rise to a three level hierarchical model (Beeravolu *et al.* 2009). The value of the parameter *m* then acts as the degree of isolation from the metacommunity.

Subsequently, the intermediate pool of species under the 3L-SINM can be modelled as a sample drawn from the metacommunity under the influence of non-neutral processes or simply as an immigration-limited sample (see Jabot *et al.* 2008 for a similar approach). Moreover, as for local communities, this intermediate pool can be described analytically using a genealogical approach (for each individual of a local community) similar to the coalescent theory of population genetics (Etienne & Olff 2004). The composition of this intermediate pool is then defined as the set of ancestor species having immigrated at some point of time into the local communities. This 3L-SINM also “collapses” back to the particular case of a 2L-SINM for the case of an immigration-unlimited random sample of the panmictic metacommunity (see Fig. 1). Besides, this collapse and vice versa forms the backbone of Munoz *et al.*’s (2007, Appendix) coalescence simulation strategy of multiple local community samples which is briefly described later (see “Simulating multiple local community samples”).

### Conditional similarity under the 3L-SINM

Following Munoz *et al.* (2008), we refer to the similarity of individuals belonging to a same species within a local community sample (*F_intra_*) using the Simpson concentration (Simpson 1949). Let *S* be the total number of species among *N* local community samples. For a given sample *k* (varying from 1 to *N*), this index represents the probability of randomly drawing (without replacement) two conspecific individuals (using Munoz *et al.*’s (2008) exact estimator of sample similarity):

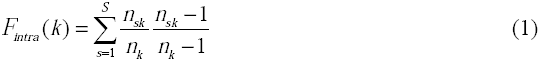

where 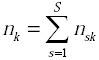 and *n_sk_* is the number of individuals of the *s^th^* species found in the *k^th^* sample. We define the time dependant version, *F*_*intra*_(*k*,*t*), of the intra sample similarity (or conspecific at time *t*) and in keeping with the 3L-SINM framework, the theoretical similarity at *t* between the intermediate (regional) pool of available immigrating species and a given local community *k* is *F*_*inter*_(*pool*,*k*,*t*). If each local sample is assumed to reasonably approximate the composition of its respective local community, it is possible to derive, as shown in the following, the time independent expectation *F_intra_*(*k*) (Munoz *et al.* 2008, Appendix C).

Let us assume that, between a time step *t* and *t*+1, there occurs a random death in the *k^th^* local community, such that two randomly chosen individuals may or may not contain the dead individual with probability 2/*n*_*k*_ and 1 − 2/*n*_*k*_ respectively. Following a coalescent approach (see Etienne & Olff 2004, eqn 1), for every dead individual, the replacing individual is either the offspring of a local individual or an immigrant individual of a lineage currently not present in the community. In the former, the replacement probability is given by *m*_*k*_ and 1 − *m*_*k*_ for the latter, where *m*_*k*_ represents the immigration probability into the *k^th^* local community. Consequently, the conspecific probability (hereafter denoted CnP) at *t*+1 of two randomly chosen individuals from a community which include the dead individual is the sum of three different transition probabilities.

When the replacement is an immigrating replacing individual, the CnP of the chosen couple is *F*_*inter*_(*pool*,*k*,*t*) which is the time conditional version of *F*_*inter*_(*pool*,*k*). If the replacement is a local event, it could be the descendant of the other individual, with probability 1/*n_k_*, in which case the CnP would be 1. If the replacing individual belongs to an offspring of an individual from the rest of the community (with probability 1 − 1/*n*_*k*_) the CnP is given by *F*_*intra*_(*k*,*t*). We could also consider the possibility that the dying individual produces offspring (a modification of Moran’s (1958) model), thus adding to the competition for the vacant spot as suggested by Hubbell (1979; 2001), though this wouldn’t affect the final result (Munoz *et al.* 2008). In sum, considering all the transition probabilities defined above, we can write the full CnP at *t*+1 for any two individuals in a local community as (Munoz *et al.* 2008, Appendix C)

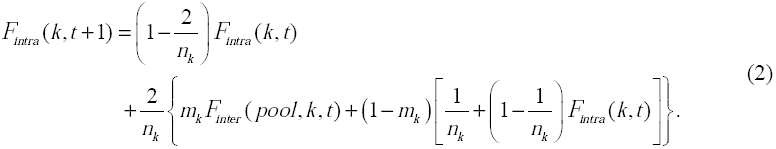

Here, *m*_*k*_ can be defined either as *I*_*k*_/(*I*_*k*_ + *n*_*k*_) or *I*_*k*_/(*I*_*k*_ + *n*_*k*_ − 1) which corresponds to the unmodified and modified Moran’s models respectively and *I*_*k*_ stands for the number of immigrating individuals into *k*^*h*^ local community (Etienne & Olff 2004). At steady state, *F*_*intra*_(*k*,*t*+*1*) = *F*_*intra*_(*k*,*t*) = *F*_*intra*_(*k*), which reduces eqn (2) to

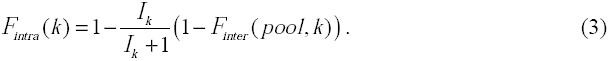

Let us assume that the *k* local community samples are far enough from each other in order to represent distinct communities and denote *F*_*inter*_(*k*) to be the CnP of an individual from the *k*^*th*^ community and an individual from one of the *k* − *1* other communities. Consequently, the coalescence approach dictates that *F*_*inter*_(*k*) is equal to the CnP of their respective ancestors who are distinct immigrating individuals from the regional pool (i.e. *F*_*intra*_(*pool*)) which can be written down as (Munoz *et al.* 2008, eqn 3):

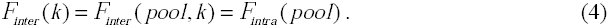

### Local similarity under the 2L-SINM

In this section, we develop the other key idea which pertains to the model “collapse” from the 3L-SINM to the 2L-SINM (see Fig. 1.) which entails that the results of the previous section also apply to the particular case of the 2L-SINM. For the sake of completeness, we also detail the analytical relationship linking *F*_*intra*_(*k*) to the immigration parameter *I*_*k*_ and the biodiversity parameter *θ* for the 2L-SINM (see also Etienne 2005, eqn 8).

Let *F*_*n*_(*k*) be the CnP of drawing a sample of *n* individuals from the *k^th^* local community and *j* the corresponding number of ancestors in the metacommunity for the same sample. In other words, the quantity *j* corresponds to the number of those individuals who were the first of every lineage to have immigrated into the community sample, which, for a sample of size n, is *j* < *n* original community lineages. Accordingly, we write the n-sample CnP as

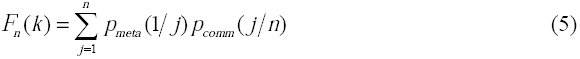

where *p*_*comm*_(*j/n*) is the probability of drawing a sample of *n* individuals from the *k*^*th*^ local community which are the progeny of exactly *j* different immigrating individuals and *p*_*meta*_(1/*j*), the probability that all of these *j* ancestors belong to the same species.

The 2L-SINM describes a metacommunity at speciation-drift equilibrium whose sample abundance distribution can be described using well known multinomial formulas from the field of population genetics (see Hubbell 2001, 119). Thus a random sample from the metacommunity containing *j* individuals belonging to σ different species has the probability

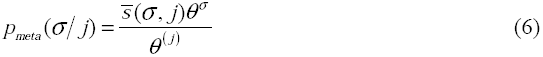

where 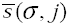 is the unsigned Stirling number of the first kind and *θ*^(*j*)^ is the Pochhammer notation for the rising factorial defined as 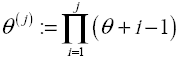 (Ewens 1972; Tavaré & Ewens 1997, eqn 41.5). This can also be extended to an immigration-limited local community (at immigration-drift equilibrium) where the immigration process replaces the speciation process and

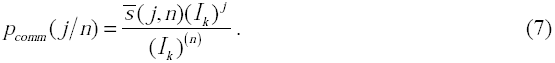

Thus, eqn (5) can be rewritten as

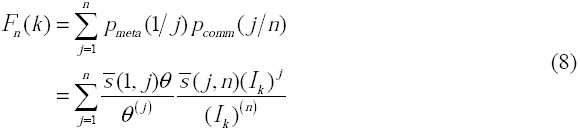

which, for the special case of a conspecific sample consisting of two individuals (i.e. *n* = 2) reduces to (Etienne 2005, eqn 8)

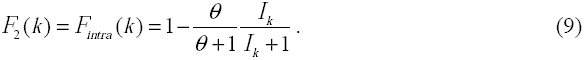

Using eqns (3), (4) and (9), the expression for *θ* simply reduces to

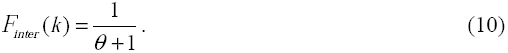

### Estimator *θ* based on the conditional inter-sample similarity

Here, we define a statistic (previously used by Munoz *et al.* 2008) which measures the *k*^*th*^ sample’s similarity with respect to the rest of the samples thereby providing an estimate of the quantity *F_inter_ (k).* This can be written in terms of sampling without replacement as

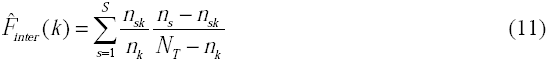

where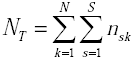 and 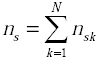. Using eqn (10), we can now write down our novel estimator for the biodiversity parameter *θ* as,

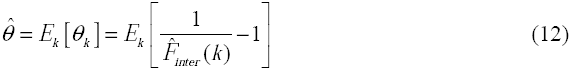

where *E*_*k*_ denotes the expectation over the *k* local community samples.

While eqn (12) can be applied directly to simulated local community samples with a known theoretical θ, it needs to be adapted for a field dataset which may belong to a single or several metacommunities. In the following we attempt to identify a subset of the field dataset which is most likely to correspond to a 2L-SINM framework (i.e. a speciation-drift equilibrium at the metacommunity level). One possible way to go about this task is to measure the spread of the 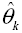 distribution and attempt to reduce it by using a sequential elimination scheme which opts out field samples one at a time.

Let us assume that our field dataset consists of *Y* community samples of variable size. Our method consists of calculating the 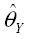 value of a given dataset along with a measure of statistical deviation (denoted COD_Y_, see below) and repeating the same by randomly pulling out one sample at a time and computing 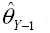 and COD_Y-1_. Among the *Y* values of 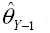 thus obtained, we eliminate the sample whose absence produced the smallest COD_*Y*−1_ value and then proceed with the sequential elimination scheme with the remaining *Y*−1 samples. We define our coefficient of deviation or COD (a robust analogue of the coefficient of variation) as the ratio of the mean absolute deviation (MAD) over the average where

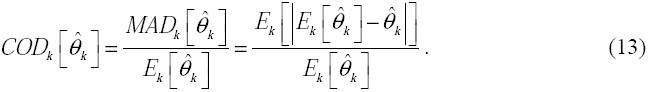

The MAD is a well known robust statistic of the sample variation when the population distribution is unknown or for highly skewed curves commonly known as heavy-tailed distributions. For a normal distribution, the standard error can be roughly calculated as 1.253 times its MAD value (MathWorks 2008). This descriptor should not be confused with the closely related median absolute deviation (also known as MAD) which uses a median instead of a mean. The elimination scheme was finally stopped with the appearance of a consistent asymptotic pattern of the sequentially obtained COD values which indicated a stable estimate for a network of samples.

## Applications

### Simulating multiple local community samples

We simulated steady state local community samples following the modification of the sequential construction scheme of Etienne (2005, Appendix S2). The simulation algorithm basically follows the 3L-SINM structure with *m* for the intermediate pool set to 1 (refer to Fig. 1.) thereby producing samples strictly congruent with the 2L-SINM. Accordingly, we create an explicit link between the ancestry information of the community samples and a large predefined metacommunity (see Munoz *et al.* 2007, Appendix).

Every simulation consists of a set of *N* local community samples, each having a randomly chosen immigration parameter *I*_*k*_ (varying from 3 to 300) and sample size *n*_*k*_ (varying from 200 to 600). Our criterion for the sample size corresponds approximately to the size of a hectare of tropical forest (i.e. *n*_*k*_ = 400 trees above 10 cm of diameter) typically found in field studies involving several permanent sampling plots (Pyke *et al.* 2001; Ramesh *et al.* 2010b). We also varied the number of samples (i.e. *N*)by simulating 5, 10, 20, 30 and 50 samples. These simulations also need a biodiversity parameter to be defined for which we simulated sets of scenarios where *θ* = (10, 50, 100, 200, 300). In the following, we shall to a simulated sampling protocol (or SSP hereafter) as a simulation generated using the information provided by the couplet (*N*, *θ*) as the other parameters are chosen to vary at random (i.e. *I*_*k*_ and *n*_*k*_). Thus, we have considered a total of 5 (values for *N*) × 5 (values for *θ*) = 25 SSPs, each of which was in turn replicated 200 times whereby we obtained a grand total of 5000 estimates (denoted 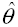;) of the theoretical biodiversity parameter θ. We assessed the performance of the estimation of *θ* by studying the histograms of the Relative Bias (RB) given by 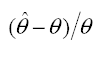 and the COD (cf. eqn (13)).

### Inferring *θ* using field data

Apart from simulations, we estimated 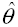 using two tropical forest datasets each consisting of multiple small permanent field plots. Both these datasets consist of the abundance information ofindividuals above 10 cm of diameter at breast height which have been identified to the species level. The first dataset consists of 50 plots (mainly 1ha) along the Panama Canal Watershed (PCW) area set up by the CTFS (Pyke *et al.* 2001), a subset of which was previously used by Jabot *et al.* (2008) for inferring immigration parameters. This freely available dataset is part of a larger study (the Marena dataset) and the field plots used in this paper are originally referred to using the following numbering 1-41, C1-C4 and S0-S4 (see Condit *et al.* 2002, Appendix; Chave *et al.* 2004, Appendix B). Our second dataset also consists of 50 field plots (each of 1ha in size) of the wet evergreen forest type from the Western Ghats (WG) region of South India (extracted from Ramesh *et al.* 2010a). This dataset has been discussed previously by Munoz *et al.* (2007, Appendix) for the purpose of inferring neutral parameters.

## Results

We studied the RB (relative bias) and the COD (coefficient of deviation, see eqn (13)) histograms for the various SSPs (simulated sampling protocols). The 5000 SSP estimates of 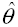 which make up the histograms were obtained in the matter of a few minutes using the MATLAB® software (MathWorks 2008). In both the five and fifty SSPs (Figs. 2 and 3 respectively), the best fit normal curve clearly emphasized the increasing symmetry of the distribution of the RB of 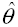 with increasing theoretical *θ* and suggested that the estimator is unbiased. In general, the RB distribution of 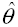 showed a tendency to be skewed for low values of *θ* while it became more symmetric and always remained centred around zero as *θ* increased (see also Figure S1 in the Supporting Information). At the same time, the COD distribution was skewed for a low number of samples (e.g. *N* = 5, 10) and a high theoretical *θ* and vice versa for *N* > 10, while the COD skewness varied little (for low *θ*) compared to the RB skewness (Figure S1). However, note that the COD histograms (Figs. 2 and 3) rarely exceed a maximum value of 0.2, which was used as a benchmark when estimating *θ* on field data for large *N*.

**Figure 2.**
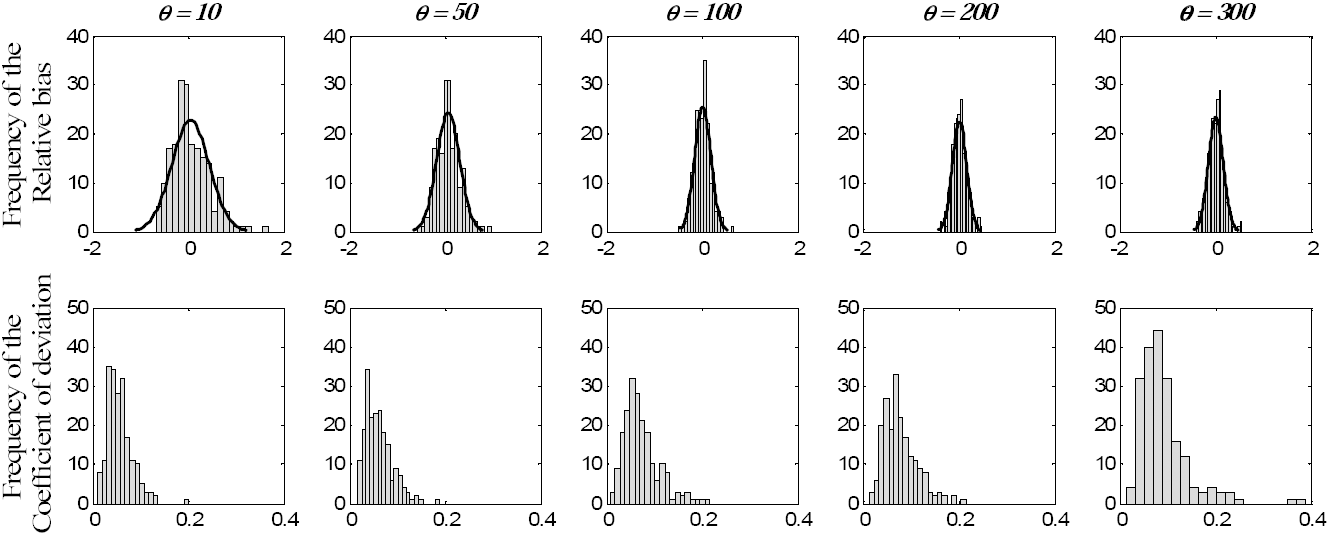
The distribution of the relative bias and the coefficient of deviation values of 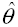(the estimated biodiversity parameter) for a sampling protocol containing 5 simulated plots each and for five different theoretical values of *θ* found in the literature. The solid black line is the best fit normal curve and emphasizes the increasing symmetry of the distribution of the relative bias with increasing *θ*.

**Figure 3.**
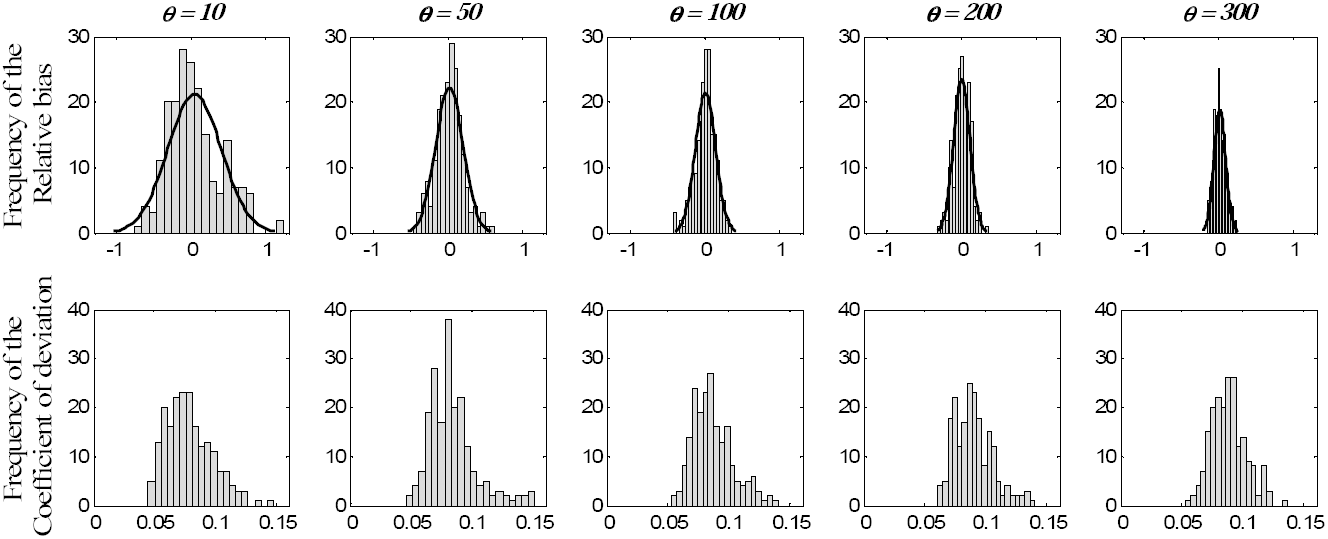
The distribution of the relative bias and the coefficient of deviation values of 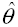 (the estimated biodiversity parameter) for a sampling protocol containing 50 simulated plots each and for five different theoretical values of *θ* found in the literature. The solid black line is the best fit normal curve and emphasizes the increasing symmetry of the distribution of the relative bias with increasing *θ*.

**Figure 4.**
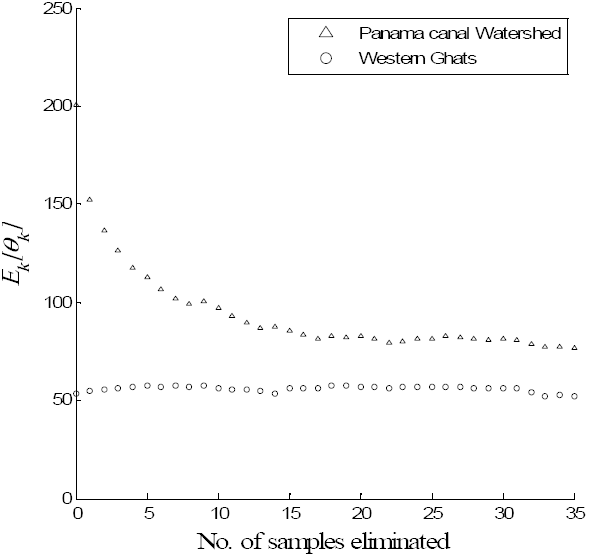
Figure 4 Estimating 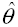 from field data using eqn (13) and following a sequential elimination scheme (see main text) in order to identify the network of plots composing the ideal 2L-SINM metacommunity. The sample elimination criterion is determined by the maximum decrease in the coefficient of deviation due to the absence of a particular plot. Here we sequentially eliminated 35 samples which appeared sufficient to highlightthe asymptotic pattern which indicated the stable 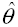 estimate for a network of plots.

We also applied our estimator to the two tropical forest datasets presented above (Fig. 4). When using all the 50 field samples of the PCW data, we obtained a high COD_50_ value (≈ 0.8) in comparison to the COD histogram for the *N* = 50 strictly neutral SSPs (Fig. 3). In contrast, the COD_50_ for the WG data (≈ 0.2) was well within the range (not shown). Subsequently, we applied the sequential elimination scheme on both these datasets (cf. previous section) in order to identify the network of plots composing the ideal (i.e. panmictic) 2L-SINM metacommunity. The estimation of the neutral biodiversity parameter for the WG dataset proved to be comparatively stable while its respective COD values fluctuated slightly (between 0.1 – 0.2) below the maximum value observed for strictly neutral simulations. Furthermore, our estimation of 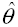 for the WG data was well bounded by the values 62.33 and 50.99 which are those found by Munoz *et al.* (2007) for the very same plots. For the PCW dataset, 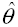 estimates reached a qualitatively stable estimate which corresponded to a COD ˂ 0.2. This was obtained after having sequentially eliminated 15 plots (COD ˂ 0.2), though we continued to eliminate samples in order to check for its stability (Fig. 4). Besides, a closer look at the remaining PCW plots revealed that the first eight eliminated plots were part of the Outer PCW region (numbered 31-39, see Pyke *et al.* 2001, Fig. 1).

## Discussion

In this paper, we have basically introduced a new estimator for multiple field samples of the neutral biodiversity parameter *θ*, first formulated by Hubbell. Subsequently, this estimator has been tested on wide-ranging simulations of multiple neutral local community samples at migration-drift equilibrium. A general conclusion from our simulation study is that the relative bias of 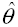 seems to be relatively well distributed around zero with a progressive tightening of the same as *θ* increased. This property is highly desired given that currently available estimators of *θ* for a single local community sample present an increasing bias with increasing *θ* (Munoz *et al.* 2007). As for the estimation variance, measured using the COD values, we note that the number of community samples used (i.e. *N*) is an important factor as a fewer number (e.g. 5 and 10) leads to an increase in the spread of the COD distribution (Fig. 2). Our results can also be seen as a significant improvement in contrast to likelihood approaches (Etienne 2009b) which become computationally intractable for more than five samples of small size like the ones used in this paper (Beeravolu *et al. Submitted manuscript*).

However, the technique presented here needs to be further compared to existing methods (Munoz *et al.* 2007; Etienne 2009a) that estimate *θ* from a single panmictic sample and also make use of the analytical expectations for the Ewens Multivariate Distribution (Ewens 1972) of population genetics literature. In particular, by randomly sampling an individual from several spatially separated samples these authors (Munoz *et al.* 2007; Etienne 2009a) constitute a metacommunity sample which is repeated a number of times in order to hone their estimates of 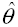. But this approach can be unreliable when the number of local community samples is small as it would provide a (relatively small) metacommunity sample with insufficient information for inferential purposes. Though, an added advantage of using the Ewens estimation for *θ* is the availability of an analytical expression for the induced bias which decreases with increasing sample size and increases with an increase in *θ* (Donnelly & Tavaré 1995, 414; Tavaré & Ewens 1997, 236).

Moreover, as our estimates using the WG data seem to be relatively robust to the sequential elimination scheme and coherent with Munoz *et al.* ’s (2007) estimates, the 50-sample evergreen forest dataset from the WG seems to strongly corroborate a 2L-SINM at this particular sampling scale. At the same time, Ramesh *et al.* (2010b) have found that some environmental variables had a strong predictive power on the plots’ floristic composition. This implies that while the data seem to agree with the neutral model, violations from neutral assumptions might not hinder a sound estimation of 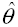 as a phenomenological descriptor of the overall diversity of the region. Instead, estimating 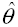 may be a good basis to compare the overall diversity between biogeographic regions as Fisher’s *α* is known to be asymptotically identical to *θ* (Hubbell 2001, 165). This approach could also be extended to other forest types and biogeographic regions in order to identify the extents of the different metacommunities as discussed above and measure their relative diversity.

To pursue this idea further, we can use the distribution of the 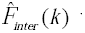 values in order to identify the local communities whose top-down linkage to a common metacommunity may be unlikely. Conversely, it is also a simple approach which delineates a subset of samples that look “floristically homogeneous” with respect to the 2L-SINM. Such a group of field plots whose taxonomic composition complies with the 2L-SINM of a metacommunity could be used to spatially delimit or map forest types over a regional sampling design. While the elimination technique presented in this paper can be seen as a simple top-down clustering scheme, it produces results which bear a close resemblance to well known bottom-up ordination schemes such as hierarchical agglomerative clustering. For example, Chust *et al.* (2006) used spatially explicit information in relation to a field plot’s ecology such as its elevation, remotely sensed data and the geographical distance to predict the forest types for almost all of the PCW data used in this paper. They perform a hierarchical agglomerative clustering of the sample abundances (or occurrences) using a proportional-link linkage algorithm and further extrapolate each cluster’s spatially explicit characteristics using a multiple regression model thereby mapping a forest type. The most “distant” clusters found in their study (Chust *et al.* 2006, Appendix 2) correspond perfectly to some of the first few plots eliminated using our sequential elimination technique. Although, note that Chust *et al.* (2006, Figs. 1 and 4) exclude some of the plots used in this paper (numbered 38 and 39) and instead use other plots (numbered P1, P2, G1, G2 and SH) which were absent from the present (see Chave *et al.* 2004, Appendix B for a corrected account of the PCW field plot numbering).

For the case of a single large local community sample, Jabot & Chave (2009), echoing Harte (2003), contend that species abundances contain a limited amount of useful information and supplement it by the use of species phylogenetic information in order to resolve inconsistencies in theestimation of neutral parameters. Our results set in a more complex context raise the question whether species phylogenies are truly needed when multiple spatially separated field samples are available. Nevertheless, the *F*_*inter*_(*k*) statistic (eqn (11)) presented here can easily be developed into an efficient Bayesian framework *(sensu* Jabot & Chave 2009), which remains a very powerful method, for injecting additional information such as species phylogenies or even the demographic history of the communities (Beaumont & Rannala 2004). Moreover, recent developments in the field of theoretical population genetics make use of a comparable inter-sample similarity metric (Gaggiotti & Foll 2010) under the island model of Wright (1931), which is a classic model of population subdivision. Interestingly, note that the island model comes conceptually close to the mainland island model of MacArthur & Wilson (1967) for the case of an infinite number of islands, in which case it is known as the continent-island model (Wilkinson-Herbots 1998, 574) or an island- mainland metapopulation (Rannala & Hartigan 1995) although there are subtle differences to be taken into account (in terms of demographic assumptions).

Finally a major weakness of almost all neutral approaches is that it is an equilibrium theory which nevertheless has greatly facilitated its mathematical development. Though, truly dynamic neutral modes are desperately lacking in community ecology (but see Leigh *et al.* 1993; Gilbert *et al.* 2006) and some initial steps have been taken in this direction (Vanpeteghem & Haegeman 2010), much needs to be done before we are able to infer the parameters of a dynamic model from field data. However, the main improvement presented in our paper is a simple and computationally efficient approach for estimating the biodiversity parameter *θ* (in the case of a multiple sample 2L-SINM framework) which is in many ways complementary to the estimation of the multiple sample *I*_*k*_ parameter (Munoz *et al.* 2008).

## Author contributions

C.B.R., P.C. and F.M. designed the study; C.B.R. performed the research; and C.B.R., P.C. and F.M. wrote the manuscript. The authors declare no conflict of interest.

## Acknowledgements

We thank Raphaël Pélissier, Olivier Hardy and Tim Keitt for helpful suggestions on a previous draft. C.B.R. is extremely grateful for the unwavering support and funding from Aurélie Loiseau towards the end of his Ph.D.

